# An R package for ensemble learning stacking

**DOI:** 10.1101/2023.06.06.543970

**Authors:** Taichi Nukui, Akio Onogi

## Abstract

**Summary:** We developed an R package for stacking, which is an ensemble approach to supervised learning. Using this package, training and prediction of stacking can be conducted using one-row scripts.

**Availability and implementation:** The R package stacking is available at the GitHub (https://github.com/Onogi/stacking).

**Supplementary information:** This manuscript has no supplementary information.

## Text

Ensemble learning is a promising approach to prediction tasks in biology (e.g., Sammut *et al*., 2022). Stacking is an ensemble learning method that can be applied to various biological processes. One application is genomic prediction, a statistical technique for predicting phenotypes/genetic merits using genome-wide DNA markers. Liang *et al*. (2021) showed that stacking using support vector regression, kernel ridge regression, and elastic nets as base learners had, on average, 7.70% higher prediction accuracy in three datasets than single models. However, stacking requires cumbersome scripts. It also requires a longer computation time to train the models because multiple models should be trained. We developed the R package “stacking” in this study to overcome these problems. This package is based on the R package caret (Kuhn, 2008), which enables users to use various supervised learning methods in the same manner via the wrapper functions provided by the package. Users can choose any model supported by care and conduct stacking without cumbersome scripting. This package implements parallel computations using the R package parallel to enable the parallel training of multiple learners.

The stacking strategy implemented in our package is illustrated below using an example of regression tasks in which the number of cross-validation (CV) folds is five, and the number of learners (members of the ensemble) is three. First, a CV is conducted with each learner using the training data. Subsequently, a meta-learner is trained using the predicted values of each learner as explanatory variables (i.e., using three explanatory variables). Because each learner is trained five times in this training process, 15 (3 × 5) base models are built. The testing data is first provided to the base models in prediction, resulting in 15 predicted values. The predicted values are averaged for each learner, resulting in three predicted values. Using these values as explanatory variables for the meta-model, the final predicted values are obtained. In classification tasks, the predicted values of the base models are not averaged. Instead, the most frequent categories are used as explanatory variables in the meta-model.

Package stacking primarily comprises four functions: *stacking_train, train_basemodel, train_metamodel*, and *stacking_predict. stacking_train* and *stacking_predict* are functions for training and prediction, respectively. The former internally calls *train_basemodel* and *train_metamodel*, but users can also call these functions themselves to optimize the base and meta-learners. Because the training process of base learners can require a long computational time, *train_basemodel* implements parallel computation using the parallel package.

*stacking_train* takes six arguments: *Y, X, Nfold, Method, Metamodel*, and *core. Y* and *X* are the vector and matrix of the objective and explanatory variables, respectively. *Nfold* is a scalar indicating the number of CV folds. *Method* is a list containing data frames as elements. The names of the elements specify the methods of the base learners and are passed on to the caret functions. Each element (i.e., the data frame) includes the hyperparameters of each learner. When the number of rows of the data frame is > 1, that is, when multiple hyperparameter values are given, all combinations of hyperparameter values are automatically created and treated as different base learners. *Metamodel* is a character that indicates the meta-learner. *core* is a scalar that indicates the number of cores required for parallel computing. *stacking_train* passes arguments *Y, X, Nfold, Method*, and *core* to *train_basemodel*; then, the list output by *train_basemodel* and argument *Metamodel* is given to *train_metamodel. train_metamodel* also takes an additional argument, *which_to_use*, that indicates which base models are used for training the meta-model. *train_metamodel* then outputs the training results of the meta-model as a list, which is then output by *stacking_train. stacking_predict* function takes two arguments: *newX* and *stacking_train_result. newX* is a matrix containing the explanatory variables of the new data, and *stacking_train_result* is the list output by *stacking_train. stacking_predict* outputs a vector of the predicted values.

First, we demonstrate the use of stacking using simulations. An explanatory variable matrix (*X*) with 1000 samples and 200 explanatory variables was randomly generated from a standard normal distribution as the training data. A vector of objective variables (*Y*) was then created by setting the regression coefficient of the first 20 explanatory variables to one and the remaining variables to zero. Interactions between neighboring variables (i.e., between 1^st^ and 2^nd^, 2^nd^ and 3^rd^, and so on) were also considered by multiplying the corresponding variables and giving regression coefficients of one. Random noise was then added such that the signal-to-noise ratio was four. Testing data of the same size were similarly generated in the same manner. As the base learners, major supervised learning methods including random forests implemented by ranger ver. 0.14.1 or 0.15.1 (Wright and Ziegler, 2017), boosting by xgboost ver. 1.7.5.1 (Chen and Guestrin, 2016) and gbm ver. 2.1.8.1 (Greg, 2007), support vector machine with a radial basis function (RBF) kernel by kernlab ver. 0.9-31 or 0.9-32 (Karatzoglou *et al*., 2004), elastic net by glmnet ver. 4.1-6 or 4.1-7 (Friedman *et al*., 2010), and partial least squares regression by pls ver. 2.8-1 (Mevik and Wehrens, 2015) were used. We used caret ver. 6.0-93 or 6.0-94 on R ver. 4.2.2 or 4.2.3, respectively. The numbers of hyperparameter sets were nine for ranger, eight for xgboost, nine for gbm, nine for support vector machine, six for glmnet, and seven for pls, resulting in 48 base learners. The hyperparameters to be specified can be confirmed using the *modelLookup* function of caret, and plausible values can be confirmed by applying methods to the data with caret. As the meta-model, elastic net implemented by glmnet was used. The training and prediction of stacking can be exceeded, for example, using the script:

stacking_predict(X.test, stacking_train(X.train, Y.train, 5, Method, Metamodel, 4)) where *X*.*test* is the explanatory variable for the testing data, *X*.*train* and *Y*.*train* are the explanatory and objective variables of the training data, respectively, 5 is the number of CV, *Method* is the list specifying the base learners and their hyperparameters, *Meta-model* specifies the meta-learner, and 4 is the number of cores used for parallel computation. The prediction accuracy, which was evaluated as the Pearson correlation coefficient between the predicted and true values, is shown in Figure 1(A). Here, stacking was compared with the methods used as base learners. Stacking showed the best accuracy on average.

**Figure 1.**
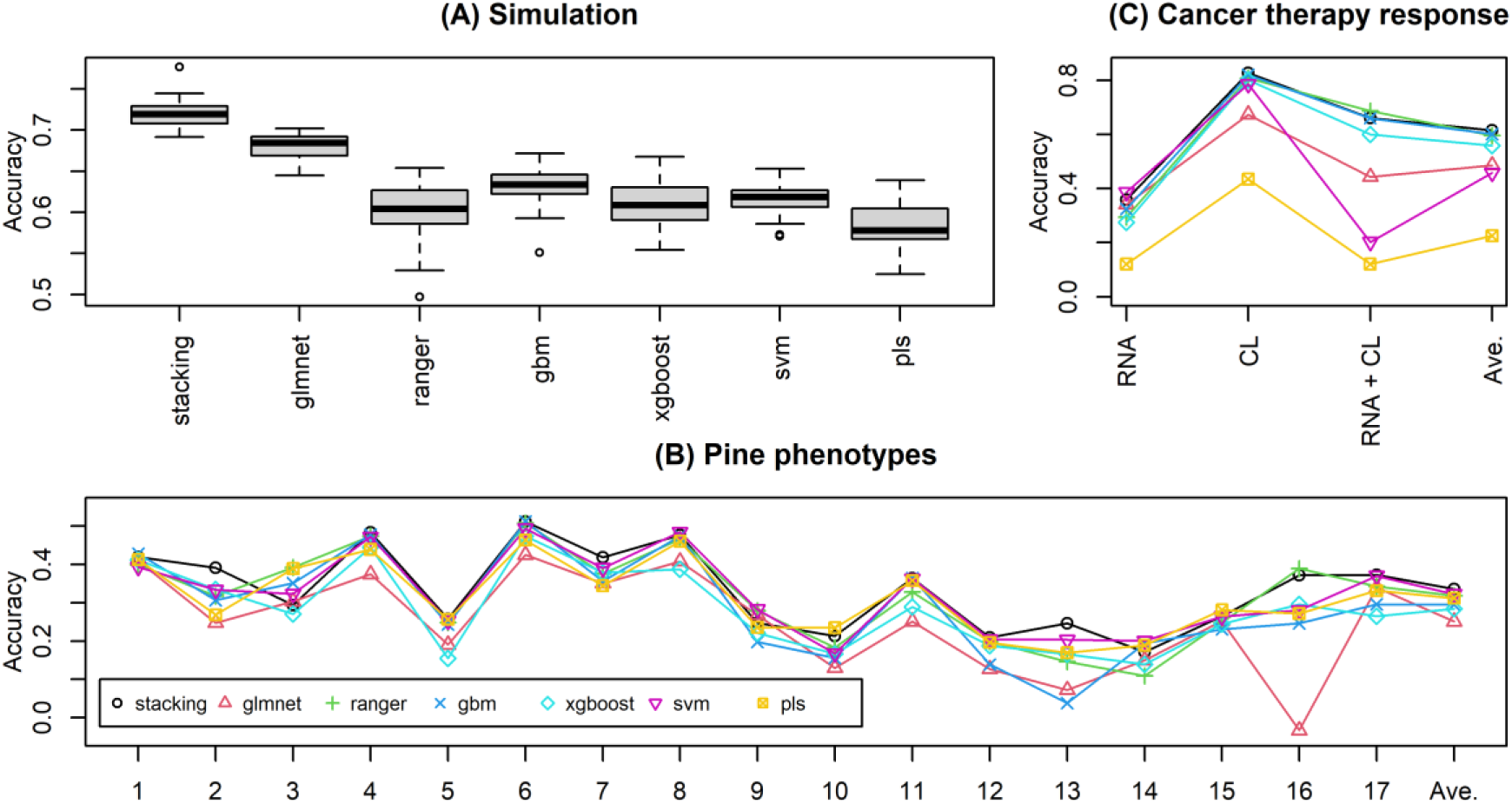
(A) Prediction accuracy evaluated with the Pearson correlation coefficients in simulation analyses (the number of replicates was 20). (B) Prediction accuracy was evaluated with the Pearson correlation coefficients in the loblolly pine data analyses. The numbers on the x-axis indicate traits. The trait abbreviations are 1, HTLC; 2, BA; 3, BD; 4, BLC; 5, C5C6; 6, CWAS; 7, CWAL; 8, DBH; 9, Density; 10, Gall; 11, HT; 12, LateWood.4; 13, Lignin; 14, Rootnum; 15, Rootnumbin; 16, Rustbin; and 17, StiffnessTree. See Resende *et al*. (2012) for the descriptions of these traits. (C) Prediction accuracy evaluated with the kappa coefficients in the cancer therapy data analyses. CL and RNA denote clinical and RNA features, respectively.

Next, we used data from loblolly pine (Resende *et al*., 2012) as an example of a regression task. The explanatory variables were the genotypes of 4853 SNPs, and the objective variables were the phenotypes of 17 traits. The number of samples was 926: 20% of samples were randomly assigned to testing data, and the remaining 80% were used for training. The base learners and hyperparameters of stacking were the same as those in the simulation study, except for *mtry* of ranger and *ncomp* of pls, which were modified according to the data size. The meta-model was also the same as that in the simulation study (elastic net). The prediction accuracy, evaluated using the Pearson correlation coefficient, is shown in Figure 1(B), where stacking exhibits the best accuracy on average. Stacking showed the best or nearly the best accuracy for most traits, whereas the rankings of the compared methods fluctuated across traits.

We then used data from breast cancer therapy as an example of a classification task (Sammut *et al*., 2022). The objective variable was a response to neoadjuvant treatment, which consisted of four categories: pCR, RCB-I, RCB-II, and RCB-III. pCR denotes no tumor, and the remaining indicates tumor residuals, in which the magnitude worsens as the number increases. To predict the response, multiple types of information, such as clinical features (e.g., tumor grade), RNA features (amounts of transcripts), and DNA features (e.g., mutation burden), were collected from the patients. Here, clinical and RNA features were used as explanatory variables because of their significant contribution to the response. Before evaluation, the top 2000 transcripts from 57905 transcripts were selected based on p-values obtained with Wilcoxon rank-sum tests between pCR and other categories to reduce computational time. The base learners used for the regression example above were also used, except for the RBF kernel of support vector machine was replaced with a polynomial kernel because the RBF kernel often failed to predict minor categories. Random forests implemented by ranger were chosen as the meta-learner. The prediction accuracy was evaluated using a five-fold CV. The kappa coefficients between the observed and predicted categories are shown in Figure 1(C), where stacking also showed the best accuracy on average and stable performance in terms of ranking among the methods, irrespective of the explanatory variables used.

In conclusion, package stacking enables users to conduct stacking without cumbersome scripts. The experiments presented show that stacking can make accurate and robust predictions, irrespective of the type of task and explanatory variables.

